# Shared vulnerability for connectome alterations across psychiatric and neurological brain disorders

**DOI:** 10.1101/360586

**Authors:** Siemon C. de Lange, Lianne H. Scholtens, Leonard H. van den Berg, Marco P. Boks, Marco Bozzali, Wiepke Cahn, Udo Dannlowski, Sarah Durston, Elbert Geuze, Neeltje E.M. van Haren, Manon H.J. Hillegers, Kathrin Koch, María Ángeles Jurado, Matteo Mancini, Idoia Marqués-Iturria, Susanne Meinert, Roel A. Ophoff, Tim J. Reess, Jonathan Repple, René S. Kahn, Martijn P. van den Heuvel, for the Alzheimer’s Disease Neuroimaging Initiative

## Abstract

Macroscale white matter pathways form the infrastructure for large-scale communication in the human brain, a prerequisite for healthy brain function. Conversely, disruptions in the brain’s connectivity architecture are thought to play an important role in a wide range of psychiatric and neurological brain disorders. Here we show that especially connections important for global communication and network integration are involved in a wide range of brain disorders. We report on a meta-analytic connectome study comprising in total 895 patients and 1,016 controls across twelve neurological and psychiatric disorders. We extracted disorder connectome fingerprints for each of these twelve disorders, which were then combined into a cross-disorder disconnectivity involvement map, representing the involvement of each brain pathway across brain disorders. Our findings show connections central to the brain’s infrastructure are disproportionally involved across a wide range of disorders. Connections critical for global network communication and integration display high disturbance across disorders, suggesting a general cross-disorder involvement and importance of these pathways in normal function. Taken together, our cross-disorder study suggests a convergence of disconnectivity across disorders to a partially shared disconnectivity substrate of central connections.

## Background

The macroscale connectome is the anatomical substrate for effective communication and integration of information between brain regions ^1,2^. Highly connected brain regions have a central role in this infrastructure forming a densely interconnected rich club core ^4,5^. This centralization of connectivity has been argued to provide several benefits for global neural integration ^6–8^ and with that healthy brain function ^9,10^. However, due to their central embedding in the network, hub regions and associated connections have also been suggested to be generally vulnerable to network disruption ^11^ and, as a result, disproportionally involved in a wide range of brain disorders ^12^.

Disease-associated alterations in structural and functional brain connectivity have been observed across a wide range of neurological and psychiatric disorders ^13,14^. Potentially, these disconnectivity patterns converge across disorders to the hypothesized vulnerable substrate of central connections. Such convergence is further suggested by observations that multiple neuropsychiatric disorders involve overlapping neural circuits ^15,16^, share genetic risk factors ^17–19^, and display high comorbidity ^20^ and shared brain phenotypes ^16^. However, so far, disease connectome studies have mostly been focused on single or small sets of disorders, and do not provide discriminative power to identify cross-disorder biological patterns of white matter disconnectivity ^21,22^.

Combining diffusion MRI data from studies on psychiatric and neurological disorders provides new opportunities to assess the vulnerability of central connections in the human brain. Here, we performed a cross-disorder data analysis, integrating connectivity alterations in a dataset comprising diffusion MRI data of in total 895 patients and 1,016 matched controls across twelve different brain disorders. These include eight psychiatric disorders (schizophrenia, bipolar disorder, attention deficit hyperactivity disorder, autism spectrum disorder, major depressive disorder, obesity, obsessive-compulsive disorder, posttraumatic stress disorder) and four neurological disorders (Alzheimer’s disease and its prodromal stage known as mild cognitive impairment, amyotrophic lateral sclerosis and primary lateral sclerosis). By combining disconnectivity maps of these twelve brain disorders we constructed a ‘cross-disorder involvement map’, identifying the set of white matter pathways that show involvement in multiple brain disorders. We combine this cross-disorder map with results from network analysis of the human connectome and show that connections important for neural integration are disproportionally involved across a range of disease processes.

## Results

### Cross-disorder involvement map

We examined MRI data of 2,681 patients and controls across twelve brain disorders from previously published studies and cohorts. Based on diffusion MRI data, white matter pathways were reconstructed in all subjects and combined with individual T1 data into connectome maps. Connectome maps were reconstructed according to a subdivision of the Desikan-Killiany atlas (DK-219). Results obtained using a second, different, parcellation of the Desikan-Killiany atlas (DK-114) are described below in the robustness analyses section. Quality control and patient-control matching was performed per study (see Methods) after which 895 patients and 1,016 matched controls were included for analysis. An overview of the demographics is provided in Figure 1 and Table 1. For each disorder, connectivity alterations were estimated in disconnectivity maps quantifying the differences in connectivity strength (fractional anisotropy) between patients and controls. Disconnectivity maps were constructed per dataset to ensure patients and controls were matched and aggregated into 12 disorders disconnectivity maps (Figure 2A). In each disorder, the top 15 % connections with highest disconnectivity effects were selected as disorder “involved”. A fixed number of connections was selected in each disorder to ensure equal presence of all disorders in the final cross-disorder involvement map.

**Figure 1.**
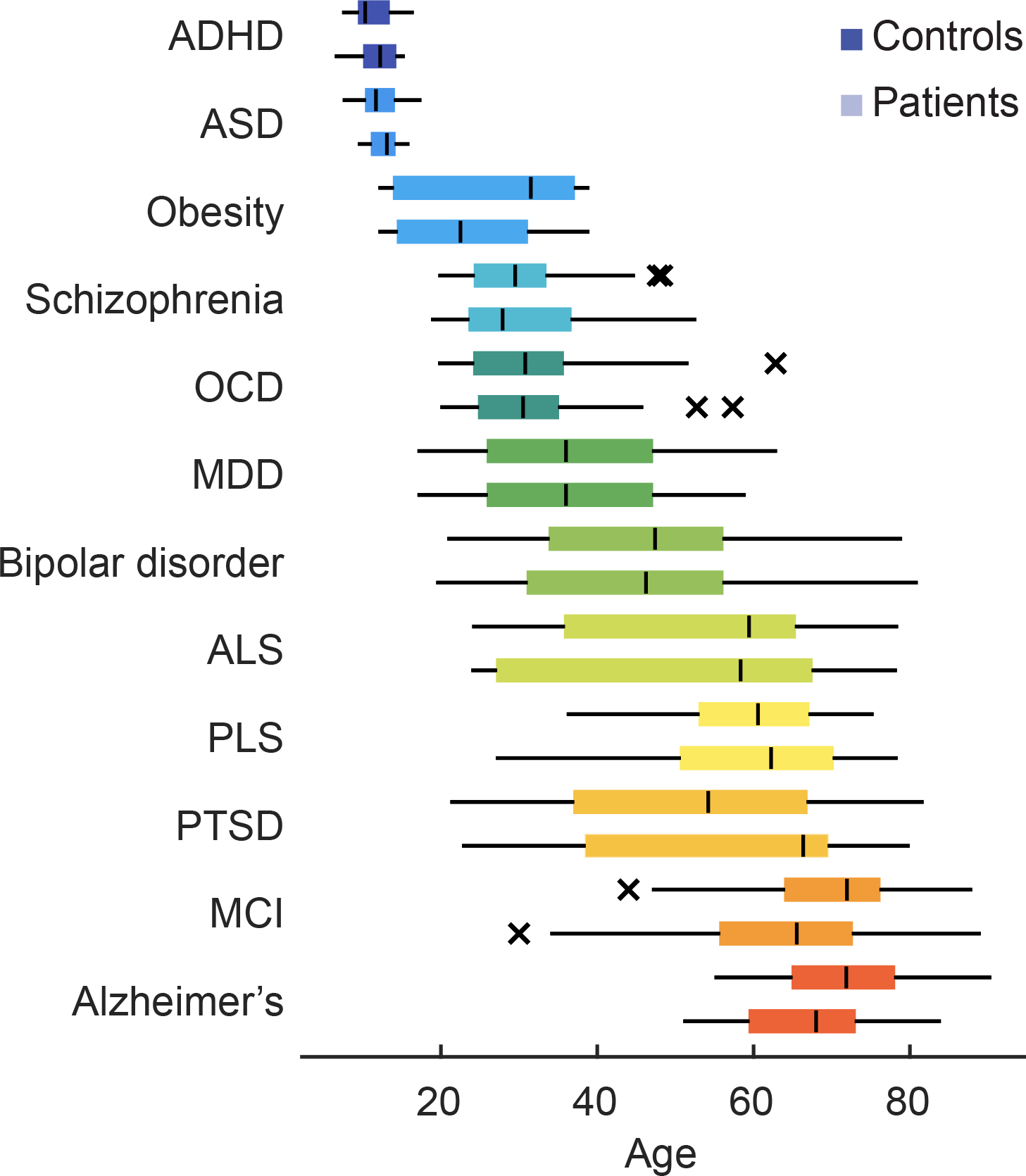
Demographics. Age distribution of controls (top) and patients (bottom) for all twelve examined disorders. Age ranged between 6 – 90 years. Controls and patients within datasets were matched on age and gender.

**Table 1.**
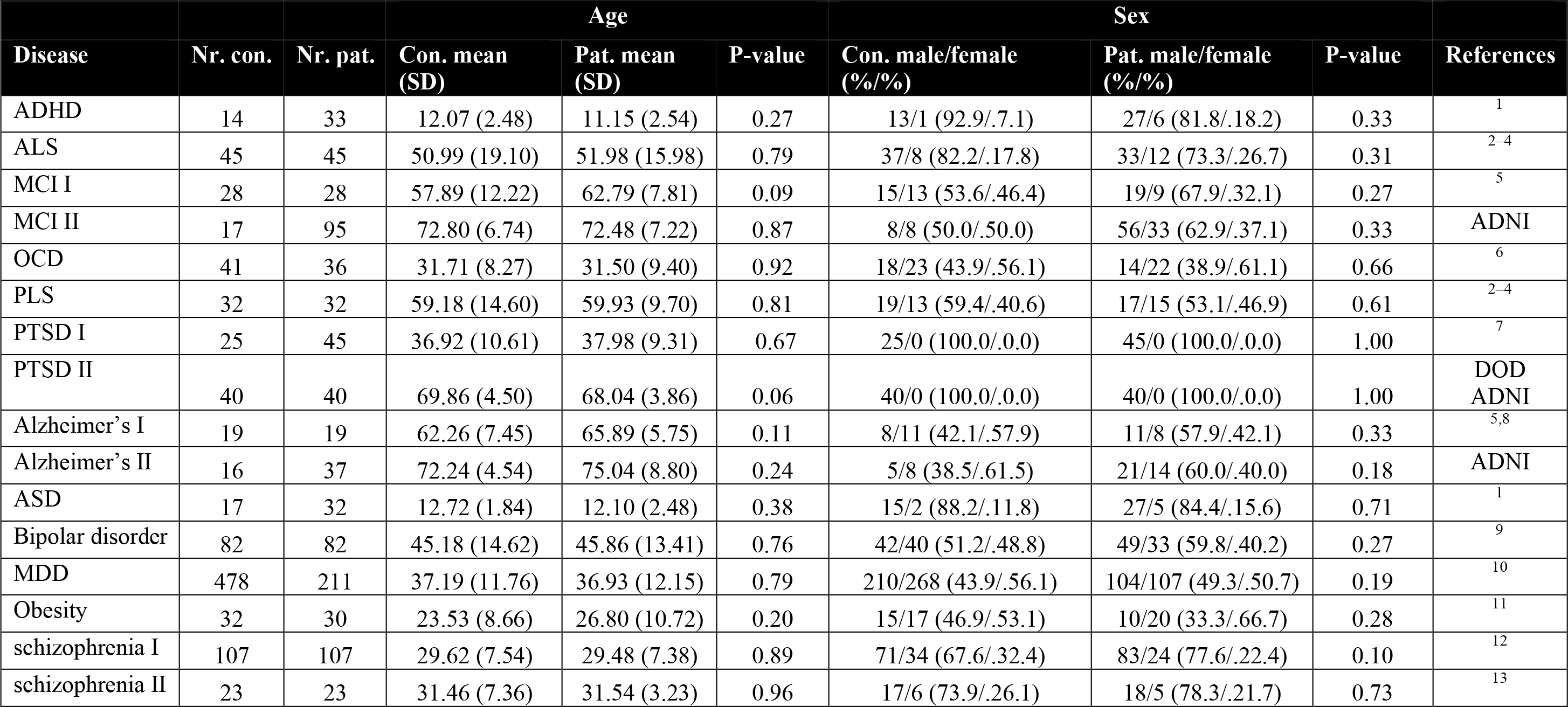
Demographics after data quality control and matching.

**Figure 2.**
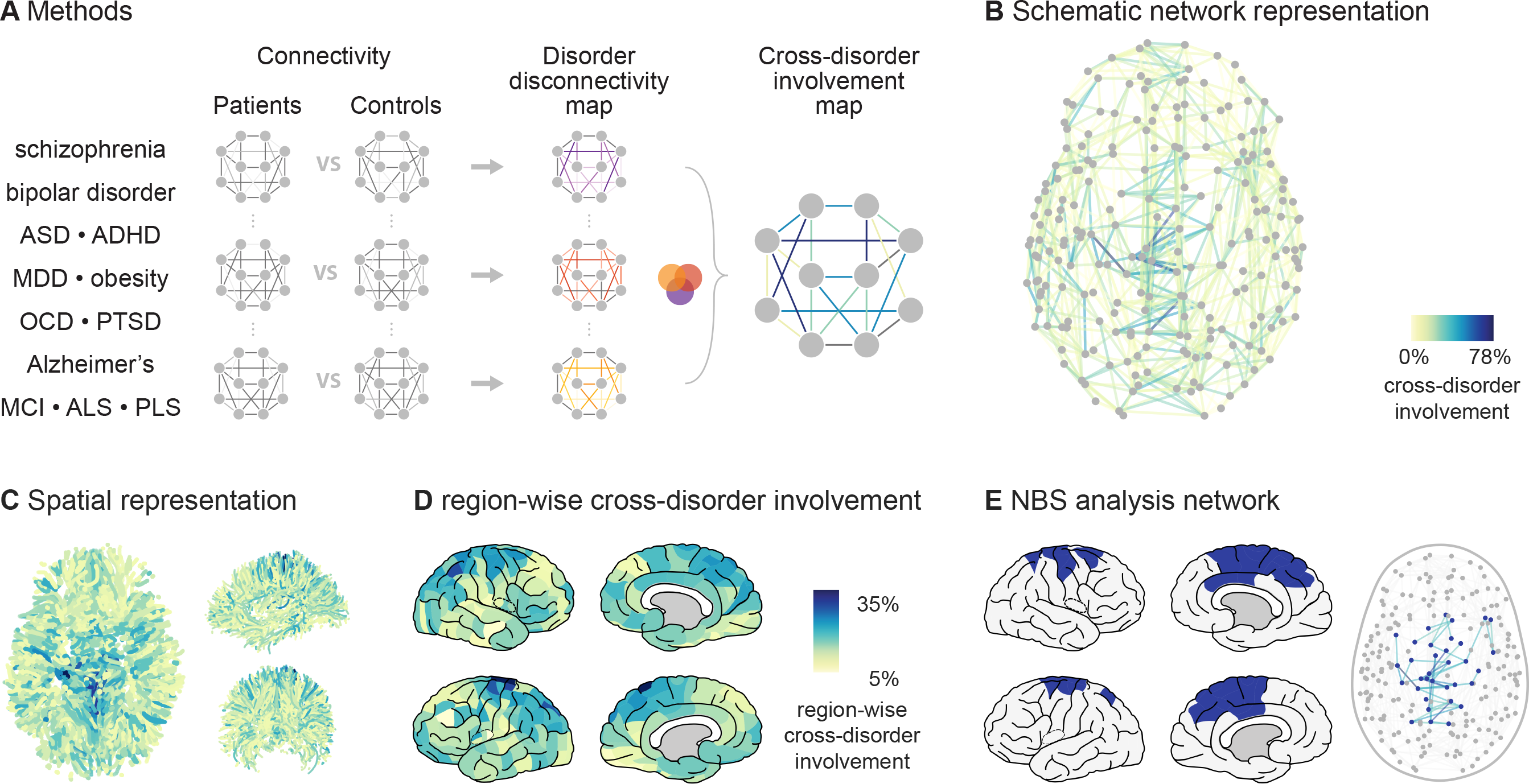
Cross-disorder involvement. **(A)**Overview of data aggregation and analysis. Per disorder, a connection-wise disorder-specific disconnectivity map was computed contrasting the fractional anisotropy of connections in patients and matched controls. Disorder-specific disconnectivity maps were combined to determine the disconnectivity distribution across disorders. **(B)** Schematic representation of human reference connectome with connections colored by cross-disorder involvement. **(C)** Superior (left panel) frontal (right-top panel) and medial (right-bottom panel) view of brain connectivity colored by cross-disorder involvement. **(D)** Lateral and medial view of left and right hemispheres showing region-wise cross-disorder involvement. **(E)** Network including 34 regions (colored blue) that showed significant involvement across disorders (NBS analysis, p = 0.0003).

Combining all individual disease maps, a group cross-disorder involvement map was then formed by the percentage of disorders in which a connection was considered “disorder involved” (Figure 2B). Network based statistics showed significantly large subnetworks of connections with cross-disorder involvement above 35%, 40% and 45% (all p < 0.05, Figure SI 1). The largest subnetwork contained of 34 regions and 82 connections, including connections of the caudal anterior cingulate, caudal middle frontal, paracentral, posterior cingulate, precentral, precuneus, superior frontal and superior parietal regions (p = 0.0003, Figure 2E). Averaging cross-disorder involvement of adjacent connections to each region provided a measure of region-wise cross-disorder involvement (Figure 2D). Regions with significantly high cross-disorder involvement included sub-regions of the postcentral gyrus (×2.23 more than in permuted cross-disorder involvement maps, p = 0.0109, FDR-corrected) and precentral gyrus (×2.19 higher, p = 0.0109, FDR-corrected).

### Network measures

#### Rich club organization

The vulnerability of the central rich club connections to disease effects was tested with respect to a rich club core of hub regions identified as the top 15% highest degree regions in the reference connectome map (degree > 14, Figure SI 2). Based on the identified rich club core, 7.6% of the network connections were classified as *rich club connections*, describing connections spanning between hub regions, 27.7% as *feeder connections*, describing connections spanning between hub and peripheral regions and 64.7% as *local connections*, describing connections between peripheral regions. Significant disproportional cross-disorder involvement was seen among rich club connections as compared to local connections (24% higher, p = 0.0041, Figure 3) and, to a lesser extent, as compared to feeder connections (17% higher, p = 0.0325). No specific increase was observed among feeder compared to local connections (p = 0.1209).

**Figure 3.**
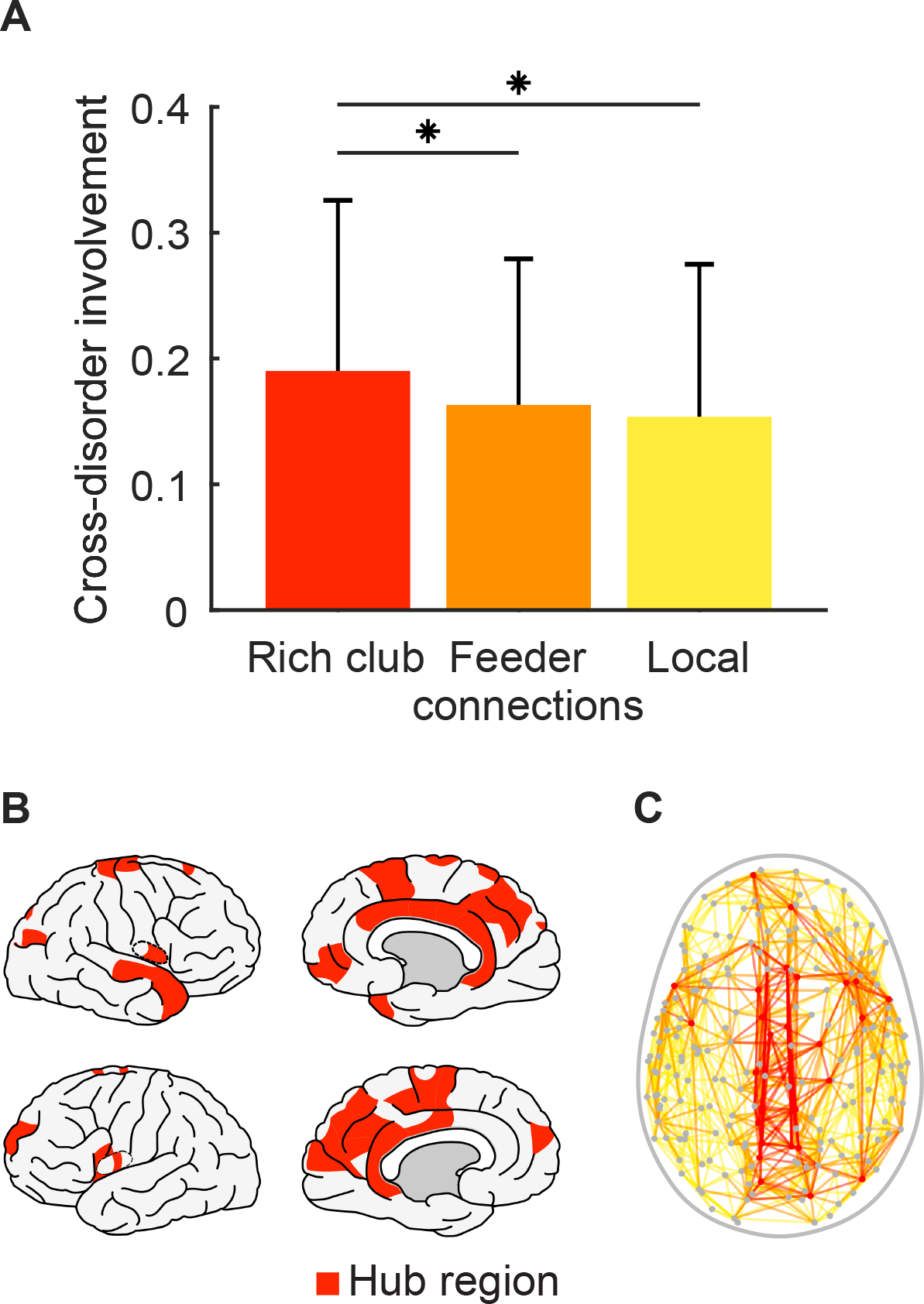
Rich club organization. **(A)** Cross-disorder involvement of rich club connections was significantly 24% higher as compared to the set of local connections (p = 0.0041) and 17% higher than observed in the set of feeder connections (p = 0.0325). Error bars mark the standard deviation. **(B)** Hub regions (top 13% highest degree regions, 29 regions) are colored in red. **(C)** Schematic representation of human reference connectome with rich club connections (colored red), feeder connections (orange) and local connections (yellow).

#### Edge-wise centrality measures

We investigated the vulnerability of central connections by examining the cross-disorder involvement of 25% most central connections identified by edge-wise centrality measures. The importance of connections for global network integration was measured by the edge-betweenness centrality, counting the number of shortest topological paths through each connection. Connections with high betweenness centrality were significantly more often involved across disorders than in randomized cross-disorder involvement maps (27% higher, p = 0.0001, Figure 4). An extended definition of global network integration is given by network communicability which considers all possible walks between nodes in the network. Connections with large edge-removal effect on the network communicability also showed significantly higher cross-disorder involvement (12% higher, p = 0.0304), further suggesting disproportionally high cross-disorder effects in connections central for global communication. In contrast, connections with strong contribution to local network organization, measured by the network clustering coefficient, did not show a predisposition for cross-disorder involvement (p = 0.7330). Finally, cross-disorder involvement was 42% increased among spatially long connections (>50 mm) in comparison with cross-disorder involvement maps with permuted disconnectivity effects (p < 0.0001).

**Figure 4.**
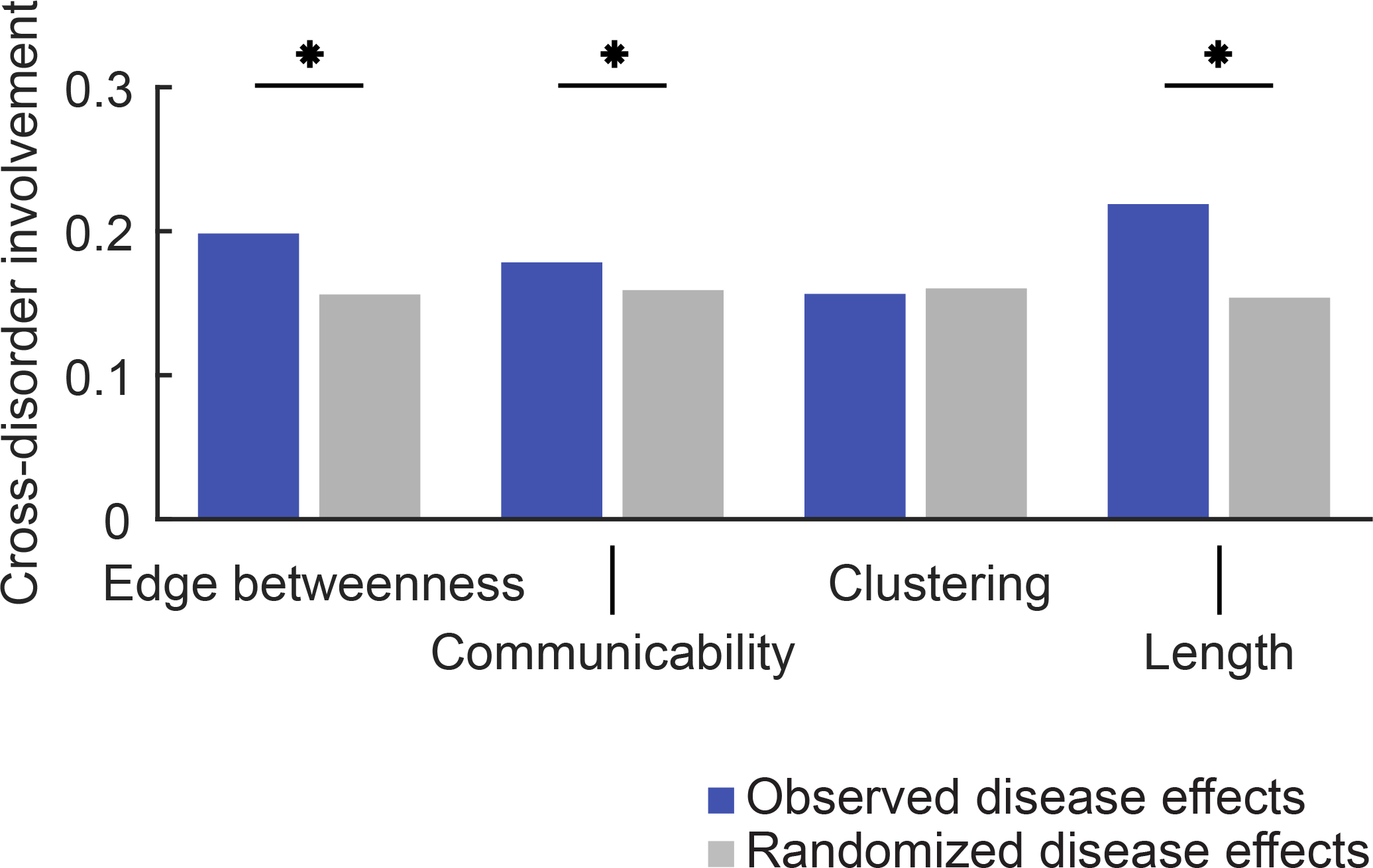
Edgewise network measures. From left to right, average cross-disorder involvement of connections with highest edge betweenness centrality (top 25% shown), highest edge-removal effect on communicability, highest edge removal effect on clustering and long-distance connections. Observed values (blue) were compared with average cross-disorder involvement in subject-label permuted maps (grey). Connections important for global topological (edge betweenness centrality, and communicability) and spatial (long-distance connections) integration showed significantly higher cross-disorder involvement levels than expected for randomly distributed disease effects (indicated by an asterisk *, p <0.05).

#### Global FA effects

To verify independence of the association between network measures and cross-disorder involvement from global FA differences (as often reported in case-control studies ^60,61^), additional permutation testing was performed in which subject labels were permuted, with now, per disease and dataset, the global FA distribution in patient and control groups preserved. Connections with high betweenness centrality again showed significantly higher cross-disorder involvement (24% increase, p < 0.0001). Connections with high edge-removal effect on network communicability also showed significantly higher cross-disorder involvement (13% increase, p = 0.0092). Cross-disorder involvement was also higher among the spatially longest connections (>50 mm), with a 36% higher cross-disorder involvement as compared to short connections (<50 mm, p < 0.0001).

#### Robustness analyses

High rich club involvement was also observed when classification of rich club, feeder and local connections were based on a smaller (7% or 9% highest degree regions) or larger set of hub regions (18% or 25% highest degree regions). For all sets of hub regions, the associated rich club connections showed significantly higher cross-disorder involvement than the associated local connections (24% - 26% higher, all p < 0.05, Figure SI 3). Also, rich club connections showed significantly higher cross-disorder involvement compared to feeder connections (14% - 22% higher, all p < 0.05, Figure SI 3) when connection classes were based on a smaller (9%) or larger set of hub regions (18% or 25%).

The results obtained using central connections selected as the top 25% connections with highest edge-wise centrality scores showed exemplarily for a range of percentages (5% - 45%) of central connections (Figure SI 4). Both smaller (10% and 15%) and larger sets (30% - 45%) of connections with high edge betweenness showed significantly higher cross-disorder involvement than seen in randomized cross-disorder involvement maps (18% - 32% higher, all p < 0.05). For central connections identified by edge-removal effect on communicability, larger sets of central connections (30% - 45%) showed significantly higher cross-disorder involvement than in randomized cross-disorder involvement maps (11% - 12% higher, all p < 0.05). Connections selected by their spatial wiring length showed at all percentages (5% - 45%) significantly higher than expected cross-disorder involvement (25% - 61% higher, all p < 0.05).

In each disorder, a fixed number of connections (15% of the connections in the reference connectome map) was selected as disorder involved to ensure equal contribution of all disorders to the cross-disorder involvement map. Validating the results obtained by selecting 15% of the connections as disorder involved, showed similar results at other reasonable percentages of disorder involved connections (5%, 10%, 20% and 25%, Figure SI 5). Rich club connections showed significantly higher cross-disorder involvement compared with local connections at all percentages (13% - 37% higher, all p < 0.05). Compared with feeder connections, rich club connections showed significantly higher cross-disorder involvement at the strict 5% (32% higher, p = 0.0241) and 10% (24% higher, p = 0.0135) percentages. Moreover, significantly increased cross-disorder involvement was observed among central connections selected by edge betweenness (18% - 42% higher, all p < 0.05) and spatial wiring length (30% - 63% higher, all p < 0.05) at all percentages. Central connections selected by edge-removal effect on communicability showed significantly higher cross-disorder involvement, when selecting the stricter set of 10% of the connections as disorder involved (15% higher, p = 0.0288).

To verify that the results were not driven by a single disorder, we performed a leave-one-out analysis in which all analyses were repeated leaving out one disorder at a time (Supplementary Materials). Results of the leave-one-out analysis showed in all iterations significantly higher cross-disorder involvement of rich club connections relative to local connections but not relative to feeder connections. The association between central connections selected by edge betweenness, edge-removal effect on communicability and spatial wiring length remained significant in all iterations (all p < 0.05, see Supplementary Materials).

Using a second parcellation atlas, we further investigated cross-disorder involvement of connections (Supplementary Materials). First, repeating the analysis revealed a subnetwork of connections with increased cross-disorder involvement similar to the network reported in the main analysis (p = 0.0002). Second, testing the vulnerability of central connections confirmed the strong findings of the main text including: significantly higher cross-disorder involvement of rich club connections compared to local connections (p = 0.0207) and strongly elevated cross-disorder involvement was observed among connections with high betweenness scores (p = 0. 0018) and spatial wiring length (p = 0.0001).

## Discussion

Our findings show that connections central to network integration and communication in the human brain are hotspots for white matter disconnectivity. Cross-disorder disconnectivity was examined in 895 patients and 1,016 matched controls across a range of twelve psychiatric and neurological disorders. Our findings provide three lines of evidence that support a general vulnerability of central connections.

First, rich club connections showed significantly higher cross-disorder involvement as compared to connections of peripheral regions (Figure 3). This observation is in line with studies showing the involvement of hub regions in specific disorders, such as schizophrenia ^62^, autism spectrum disorder ^63^, ADHD ^63^, Huntington’s disease ^64^, Alzheimer’s disease ^65–67^, and general white matter lesions ^68^. High involvement of rich club connections across disorders is further in line with studies on cross-disorder gray matter abnormalities. A large voxel-based morphometry meta-analysis showed hub regions to be disproportionally involved in anatomical abnormalities across clinical brain disorders ^12^, findings verified by voxel-based morphometry meta-analysis across clinical psychiatric disorders, showing shared gray matter loss in in particular dorsal anterior cingulate and insula hub regions ^15^.

Second, edgewise network measures revealed connections critical for network efficiency and communicability to display high cross-disorder involvement (Figure 4). This result extends earlier reported decreased efficiency of structural networks in for example depression ^65^ and in Alzheimer’s disease, schizophrenia, multiple sclerosis and ALS (see ^22^ for a review), suggesting that these effects are not disease-specific, but perhaps more general to brain disorders than previously reported. Furthermore, these results stress the hypothesized importance of efficient integration of information for healthy brain function ^3^, with disruptions in central connections potentially leading to disproportional effects in brain dysfunction.

A third line of evidence for the vulnerability of central connections is the observation of high cross-disorder involvement among connections spanning long physical distances (Figure 4). This observation is in line with studies reporting affected long fiber tracts including the superior and inferior longitudinal fasciculus in for example ADHD ^69^, ASD ^70^, OCD ^71^, and schizophrenia ^72,73^. Post-hoc analysis showed these effects to be reduced when restricting the analysis to intrahemispheric connectivity of either the left (19% increase, p = 0.0190) and right hemisphere (p = 0.0743 (n.s.)), suggesting high cross-disorder involvement of spatially long connections to be partly driven by a clustering of effects among interhemispheric connections.

The observed cross-disorder effects are likely to reflect the combination of multiple disease mechanisms that differ across disorders ^74,75^. Central regions and connections have been argued to be biologically expensive, characterized by complex neuronal architecture ^76^, high metabolism ^3^ and high neuronal activity ^77^. This high biological cost might result in increased vulnerability to a wide range of disease processes, such as a toxic environment or reductions in the supply of oxygen or other metabolic resources ^78^. Central connections may also display a high cross-disorder involvement as the result of their topological centrality and associated risk to propagating disease processes ^75,79^. Connectome studies of disconnectivity in ALS ^80^, Alzheimer’s disease ^79,81,82^ and frontal temporal dementia ^82^ have suggested a prion-type of spread of disease processes in neurodegenerative disorders with specifically early disease involvement of hub regions and rich club connections due to their central embedding in the network. In addition to this, long-range central connections may be particularly vulnerable to focal white matter degeneration. The chance of focal degeneration is proportional to fiber length, making long-range central connections in total more vulnerable to general white matter atrophy as compared to short range connections. Rich club connections have also been shown to display a prolonged development ^83–85^, which may further increase their general vulnerability by putting these connections at risk to late neurodevelopmental stress, substance use and dysregulation of hypothalamic-pituitary-adrenal axis function ^74,86^. The shared vulnerability of central connections across disorders might result from the importance of central connections for cognitive function ^4,87^. Cognitive impairment is shared across the symptomatology of many brain disorders ^88^. Hence, if in each disorder separately disconnectivity of central connections is associated with deficits in cognitive function, then such overlap in symptomatology would result in general vulnerability of central connections.

Genetics and heritability studies offer the potential to gain further understanding in the pathology underlying cross-disorder disconnectivity. Shared genetic etiology is observed across many psychiatric and neurological disorders ^17,89,90^, with shared genetic risk factors providing converging evidence for common underlying biological processes across brain disorders ^16,17,91^. Further exploring structural disconnectivity and genetic information in a multi-modal and cross-disorder approach may further identify cross-disorder as well as disorder-specific biological pathways ^16,92–94^.

The observation of overlapping disconnectivity patterns across brain disorders is in agreement with the hypothesis that brain disorders are interrelated ^17^ and prompts for a careful consideration of disease disconnectivity findings. Disconnectivity findings of single-disorder connectome examinations may often be interpreted as disorder-specific disconnectivity effects, which might not fully be the case considering the demonstrated overlap in effects across disorders. This misattribution is perhaps most problematic in the development of biomarkers for brain disorders based on disconnectivity fingerprints, where it could result in overestimation of the disorder specificity of a presented biomarker.

Methodological issues have to be considered when interpreting our findings. While combining data from multiple studies may implicitly account for real-world heterogeneity and improve generalizability of observed results ^95^, it is likely that combining data from multiple studies may also reduce statistical power as a result of inter-study heterogeneity in diagnoses, demographics, scanner and MRI acquisition protocols. We are aware of this limitation and aimed to minimize the influence of study specific properties by directly comparing control and patient data within each study first, before combining information across the twelve disorders. Second, disorder disconnectivity fingerprints were based on structural brain networks obtained by diffusion-based MRI, with white matter microstructural integrity assessed by means of the metric of fractional anisotropy ^47^. Fractional anisotropy is however only an indirect marker of the micro-scale neuroarchitecture and diffusion weighted imaging has recognized limitations with respect to the reconstruction of complex fibers and connectome mapping ^45,96,97^, which might result in underestimation of disconnectivity effects within and across disorders. Third, our conclusions are based on effects seen across twelve disorders, and it remains unclear whether our conclusions could be generalized to an even wider range of brain disorders. To verify that the results were not driven by a single disorder, we performed a leave-one-out validation analysis in which all analyses were repeated leaving out one disorder at a time. Moreover, we possibly missed smaller sets of disorders that share disconnectivity patterns. Investigating potential clustering of disorders based on their disconnectivity patterns would be of great interest to further provide new insights in more detailed biological relationships between disorders.

Our findings suggest shared connectome pathology across neurological and psychiatric disorders, with in particular high general vulnerability of connections central to neural communication and integration. Beyond identifying cross-disorder disconnectivity, cross-disorder examination has the important potential to show distinct disconnectivity patterns between disorders. Future examination into both disorder-shared and disorder-specific disconnectivity effects provides better understanding of which brain alterations are general and which effects are unique for brain disorders, providing new ways for the development of MRI based biomarkers for psychiatric and neurological disorders.

## Methods

### Studies and subjects

Diffusion MRI data of 2,681 patients and controls of twelve disorders were included. Data included diffusion-weighted imaging (DWI) data of previously reported studies on schizophrenia (two datasets available, set I and II) ^23,24^, bipolar disorder ^25^, attention deficit hyperactivity disorder (ADHD) ^26^, autism spectrum disorder (ASD) ^26^, major depressive disorder (MDD) ^27^, obesity, obsessive-compulsive disorder (OCD) ^28^, posttraumatic stress disorder (PTSD, two datasets, set I and set II) (ADNI-DOD adni.loni.usc.edu and ^29^), and four neurological disorders, Alzheimer’s disease (AD, two datasets, set I and set II) (ADNI and ^30,31^), mild cognitive impairment (MCI, two datasets, set I and set II) (ADNI and ^30,31^), amyotrophic lateral sclerosis (ALS) ^32^–34 and primary lateral sclerosis (PLS) ^32–34^. Figure 1 provides an overview of all data included and a summary is provided in Table 1. Further details including a description of MRI acquisition protocols and demographics are outlined in the Supplementary Materials. Within each dataset, patients and controls were matched on age, sex, scanner settings and where possible other demographics (procedure described in the Supplementary Materials).

### Data processing

#### DWI Tractography

Data preprocessing of DWI and T1-weighted images of individuals included the following steps: the anatomical T1-weighted image was parcellated into 219 distinct cortical regions (111 left-hemispheric and 108 right-hemispheric regions) according to a subdivision of FreeSurfer’s Desikan-Killiany atlas ^35,36^ using FreeSurfer ^37^. Using a second different parcellation of the Desikan-Killiany atlas (DK-114) showed similar results presented in the robustness analyses section and Supplemental Materials. Second, the individual parcellation map was co-registered to the DWI data using an affine transformation mapping of the T1-weighted image to the DWI image. Third, diffusion-weighted images were corrected for eddy current distortions and head motion using the FMRIB Software Library ^38^. If reversed phase encoding data was available (datasets listed in SI Table 1), susceptibility induced distortions were estimated and incorporated in the preprocessing ^39^. Fourth, a tensor was fitted to the diffusion signals in each voxel using a robust tensor fitting algorithm ^40^ and subsequently fractional anisotropy (FA) was derived ^41^. Given the mostly clinical diffusion MRI protocols used for data acquisition, simple deterministic tensor reconstruction (DTI) (as compared to more advanced diffusion profile reconstruction methods) was used to minimize the potential influence of false positives on network reconstruction and subsequent computation of network metrics ^42–44^. This relatively simple reconstruction of the diffusion signal is a limitation of our cross-disorder examination, potentially leading to incomplete reconstruction of complex fiber pathways and an underestimation of cross-disorder disease effects ^45^. Fifth, white matter pathways were reconstructed using fiber assignment by continuous tracking (FACT) ^46^, with streamline reconstruction starting from eight seeds in every cerebral white matter voxel. Fiber tracking was continued until a streamline showed high curvature (> 45º), exited the brain mask, or when a streamline entered a voxel with low fractional anisotropy (< 0.1). The mean FA value of a streamline was computed as the weighted average FA value over all voxels that a streamline passed.

#### Network reconstruction

For each individual dataset, reconstructed streamlines and cortical parcellation were combined into a weighted network. The 219 cortical areas were chosen as nodes in the network and two regions were considered connected if at least one reconstructed streamline was found to touch both cortical regions. The weight of connections was taken as the mean fractional anisotropy (FA) of streamlines involved ^47^.

### Cross-disorder analysis

Cross-disorder examination of disorder-related disconnectivity was performed in two steps. Patient and control data were *first* compared *within* each dataset (in contrast to the alternative of pooling all data into one large dataset) to ensure that patients and controls were matched on age, sex and other demographics and scanner settings. This comparison provided for each disorder a disconnectivity map quantifying the differences in connectivity strength between patients and matched controls. *Second*, disorder disconnectivity maps were combined *across* the twelve disorders to determine the distribution of disconnectivity effects across network connections of the brain. This two-step approach optimized comparability of data across studies with different MRI acquisition protocols. In what follows, we describe this procedure in more detail, including the construction of the disorder disconnectivity maps and the cross-disorder involvement map, followed by the performed statistical analyses.

### Step 1: Disorder disconnectivity map

Per disorder, a disconnectivity map was constructed by assessing the between-group difference in FA of connections between patients and controls quantified by a Student’s t-test statistic. As such, we tested for lowered FA connectivity strength in the patient group compared to the controls. Between-group analysis was performed for connections that were present in 30% or more of the population of controls and patients to ensure sufficient statistical power ^33^. To correct for possible differences in degrees of freedom across connections, t-test statistics were transformed to z-scores.

For the disorders PTSD, schizophrenia, Alzheimer’s disease and MCI, for which multiple datasets were available, a disorder disconnectivity map was first calculated *per dataset* and then combined into an average disorder disconnectivity map using Stouffer’s method for combining independent tests by averaging the z-scores in the disorder disconnectivity maps across datasets ^48,49^.

In total, this resulted in a disorder disconnectivity map for each of the 12 included brain disorders. Next, the top 15% connections with highest z-scores were selected as the set of most involved connections in that disorder, performing, per disorder, a proportional thresholding on the disorder-specific disconnectivity map with a density of 15% ^50^. Results using 5%, 10%, 20% or 25% involved connections are presented in the robustness analyses section.

### Step 2: Cross-disorder involvement map

The twelve thresholded disorder disconnectivity maps were combined into a total *cross-disorder involvement map*. To maximize comparability across studies and to avoid any potential bias to one of the included datasets, connection effects were included for those connections present in a reference group connectome map based on high-quality data of the Human Connectome Project (HCP, 500 Subjects Release of the Human Connectome Project) ^51,52^ (see Supplementary Materials for details on the HCP group connectome reconstruction). Finally, a *cross-disorder involvement map* was formed by adding up all thresholded disorder disconnectivity maps and dividing it by the number of disorders in which each connection was present, thus computing per connection the percentage of disorders in which this connection was involved.

### Network analysis

The centrality of connections in the network structure was considered with respect to rich club organization, edgewise global and local network measures and physical wiring length. Metrics were computed on the HCP group connectome to ensure independence of the examined datasets.

#### Rich club organization

Central connections were identified with respect to the rich club organization describing the collective of high-degree hub regions ^4^. Regional degree was computed on the basis of the HCP group connectome with hub regions selected as regions with a degree above 14 (top 13% regions with the highest regional degree, 29 regions, Figure 3, listed in SI Table 2). This set of regions was verified to display a rich club organization, showing a higher-than-expected level of interconnectivity (p < 0.0001, compared with 10,000 degree-preserved rewired networks using permutation testing).

Based on the identified rich club organization, network connections were classified into *rich club connections*, describing connections spanning between hub regions, *feeder connections*, describing connections spanning between hub and peripheral regions, or *local connections*, describing connections between peripheral regions ^6^. Analyses were repeated with connections classes derived from a smaller and larger set of hub regions (top 7% highest degree regions, degree > 16; top 9% highest degree regions, degree > 15; top 18% highest degree regions, degree > 13 and top 25% highest degree regions, degree > 12).

#### Global network organization

Global network integration was examined from the perspective of the ease of communication between nodes in the network. First, the centrality of connections with respect to the shortest topological paths in the network was measured by counting the number of shortest topological paths through each network connection using the metric of edge betweenness ^53^. Second, network integration was considered by examining the metric of network communicability, measuring all possible walks between nodes ^54^. The contribution of connections to communicability was assessed by edge-removal statistics ^55–57^. Removal-effect of each connection on network communicability was quantified as the difference (in terms of percentage) between the network communicability before and after removal of a connection.

#### Local network organization

The role of network connections in local network organization was assessed through the contribution of each connection to *network clustering* ^53^. The removal-effect of each connection on global network clustering was quantified as the difference (i.e., percentage of change) in global clustering before and after removal of the connection.

#### Spatial embedding

Projection length of each connection was calculated as the average physical length of a connection.

### Statistical analysis

#### Cross-disorder involvement

Significant subnetworks in the brain with increased cross-disorder involvement levels were identified using Network Based Statistics ^59^. The cross-disorder involvement map was binarized by including connections with cross-disorder involvement percentages above a specified NBS-threshold. Multiple NBS-thresholds (0%, 5%, …, 100%) were considered, capturing the trade-off between specificity and sensitivity of the NBS-analysis. The number of connections in the greatest component of the thresholded network was counted. Significance of this cluster was assessed using permutation testing by comparison with the distribution of greatest component sizes in a null condition in which disease effects were randomized. For this, for each permutation, a cross-disorder involvement map was calculated on a permuted subject sample in which subject labels (i.e. controls and patients) were randomly reassigned (keeping patient and control group sizes intact). 10,000 permutations were examined and the percentage of the permutations in which the greatest component was larger or equal to the observed greatest component was assigned as p-value to the observed cross-disorder involvement. Regions with significantly high cross-disorder involvement were similarly identified by comparison with the sample of subject-label permuted cross-disorder involvement maps. To correct for multiple testing, p-values were adjusted by the false discovery rate correction procedure ^58^.

#### Network measures

Differences in mean cross-disorder involvement between rich club and feeder, rich club and local, and feeder and local connection classes were statistically assessed using permutation testing (10,000 permutations). In each permutation, connection class labels were randomly shuffled and mean cross-disorder involvement of the classes was computed over the permuted connections. Differences in cross-disorder involvement between connection classes were computed for all permutations. The observed difference in cross-disorder involvement between two connection classes was assigned a p-value by computing the percentage of permutations in which the difference between the two connection classes was equal to or exceeded the observed difference.

The 25% connections most central connections selected by global network integration, local network integration and the spatial embedding were examined. Exploring other reasonable percentages (5%, 10%, …, 45%) for selecting central connections showed consistent results that are reported in the robustness analyses. Cross-disorder involvement levels were compared with the levels expected when disconnectivity was randomly distributed using permutation testing, this to verify independence of our results from connection properties such as connection prevalence or group-average connection strength. For each permutation, subject labels were randomly reassigned and cross-disorder involvement maps were calculated using the permuted subject-labeling. 10,000 permutations were computed and cross-disorder involvement levels of the subsets of central connections were calculated for each permutation. Based on this null distribution, the original effect was assigned a p-value as the percentage of permutations in which the cross-disorder involvement was equal to or exceeded the observed cross-disorder involvement.

#### Global FA effects

Additional permutation testing was performed to verify independence of our results from global FA differences that are often reported in case-control studies^60,61^. For each subject, global FA was computed as the total FA strength of all connections. Next, subjects were classified into ten global FA groups, group one with global FA in the interval [0, 0.1), group two with global FA in the interval [0.1, 0.2), etc. For permutation testing, subject labels were permuted within datasets, but now under the constraint of only allowing switching patient and control labels of subjects assigned to the same global FA bin. As such, the resulting global FA distribution of permuted patient and control groups was kept similar to the original global FA distributions (and therewith also potential between-group differences in global FA). 10,000 permutations were computed and, in each permutation, the cross-disorder involvement of the subsets of connections was calculated. Observed effects were assigned a p-value as the percentage of the permutations in which the measured effect was equal to or exceeded the observed effect.

## Acknowledgements

M.P. van den Heuvel was funded by an ALW open (ALWOP.179) and VIDI (452-16-015) grant from the Netherlands Organization for Scientific Research (NWO) and a Fellowship of MQ.

The Muenster Depression Cohort was funded by the German Research Foundation (DFG, grant FOR2107 DA1151/5-1 and DA1151/5-2) to UD; SFB-TRR58, Projects C09 and Z02 to UD) and the Interdisciplinary Center for Clinical Research (IZKF) of the medical faculty of Münster (grant Dan3/012/17 to UD).

Data collection and sharing for this project was funded by the Alzheimer’s Disease Neuroimaging Initiative (ADNI) (National Institutes of Health Grant U01 AG024904) and DOD ADNI (Department of Defense award number W81XWH-12-2-0012). ADNI is funded by the National Institute on Aging, the National Institute of Biomedical Imaging and Bioengineering, and through generous contributions from the following: AbbVie, Alzheimer’s Association; Alzheimer’s Drug Discovery Foundation; Araclon Biotech; BioClinica, Inc.; Biogen; Bristol-Myers Squibb Company; CereSpir, Inc.; Cogstate; Eisai Inc.; Elan Pharmaceuticals, Inc.; Eli Lilly and Company; EuroImmun; F. Hoffmann-La Roche Ltd and its affiliated company Genentech, Inc.; Fujirebio; GE Healthcare; IXICO Ltd.; Janssen Alzheimer Immunotherapy Research & Development, LLC.; Johnson & Johnson Pharmaceutical Research & Development LLC.; Lumosity; Lundbeck; Merck & Co., Inc.; Meso Scale Diagnostics, LLC.; NeuroRx Research; Neurotrack Technologies; Novartis Pharmaceuticals Corporation; Pfizer Inc.; Piramal Imaging; Servier; Takeda Pharmaceutical Company; and Transition Therapeutics. The Canadian Institutes of Health Research is providing funds to support ADNI clinical sites in Canada. Private sector contributions are facilitated by the Foundation for the National Institutes of Health (www.fnih.org). The grantee organization is the Northern California Institute for Research and Education, and the study is coordinated by the Alzheimer’s Therapeutic Research Institute at the University of Southern California. ADNI data are disseminated by the Laboratory for Neuro Imaging at the University of Southern California.

**Figure SI 1. Subnetworks identified by network based statistics. (A)** Number of regions in the greatest component in thresholded version of the cross-disorder involvement map across a range of thresholds (0% - 100% cross-disorder involvement). The greatest components ranged from including all regions (at 0% cross-disorder involvement threshold) to including only one region (at 100% cross-disorder involvement threshold). At 35%, 40% and 45% cross-disorder involvement thresholds, the identified subnetwork showed significantly larger than subnetwork seen in subject-label permuted cross-disorder involvement maps (indicated by an asterisk *, p < 0.05). **(B)** Subnetworks and included regions (in blue) of the three identified significantly large subnetworks.

**Figure SI 2. Rich club coefficient in reference connectome.** Reference connectome data showed a significant rich club organization at all degree levels above 8 (indicated by an asterisk *, p < 0.05, FDR-corrected).

**Figure SI 3. Rich club organization across percentages of hub regions.** Ratio between cross-disorder involvement of rich club and local connections (left) and feeder connections (right). The ratios were evaluated for rich club, feeder and local connections derived from sets of hub regions selected at different percentages (7%, degree > 16; 9%, degree > 15; 13%, degree > 14; 18%, degree > 13; 25%, degree > 12). Percentages at which the ratio was significantly large (i.e. significant differences in cross-disorder involvement of rich club connections and feeder or local connections) are indicated by an asterisk * (p < 0.05).

**Figure SI 4. Edgewise network measures across percentages of central connections.** The cross-disorder involvement of central connections (selected by edge betweenness (left), edge-removal effect on communicability (middle) and spatial wiring length (right)) relative to cross-disorder involvement observed in subject-label permuted cross-disorder involvement maps. The relative cross-disorder involvement was obtained at different selection percentages ranging from considering the top 5% most central connections to the top 45% most central connections. Percentages at which the ratio was significantly high (i.e. the set of central connections showed significantly higher cross-disorder involvement than in permuted cross-disorder involvement maps) are indicated by an asterisk * (p < 0.05).

**Figure SI 5. Cross-disorder involvement of central connections across percentages of disorder involved connections.** Results were computed across various percentages of connections selected as disorder involved in addition to the 15% percentage used in the main analysis. **(A)** The ratio in cross-disorder involvement between rich club and local (left) and feeder (right) connections. **(B)** The relative cross-disorder involvement of central connections compared with subject-label permuted cross-disorder involvement maps. Significant effects are indicated by an asterisk * (p < 0.05).

**SI Table 1.** Acquisition parameters of included datasets.

**SI Table 2.** Number of excluded subjects (because subjects miss information, subjects are considered outlier, or subjects are not matched) per dataset.

**SI Table 3.** List of hub regions (region subnumbers are study specific).

